# IL-7 armed binary CAR T cell strategy to augment potency against solid tumors

**DOI:** 10.1101/2025.06.25.661524

**Authors:** Alejandro G Torres Chavez, Mary K McKenna, Anmol Gupta, Neha Daga, Juan Vera, Ann M Leen, Pradip Bajgain

## Abstract

Clinical studies of T cells engineered with chimeric antigen receptor (CAR) targeting CD19 in B-cell malignancies have demonstrated that relapse due to target antigen (CD19) loss or limited CAR T cell persistence is a common occurrence. The possibility of such events is greater in solid tumors, which typically display more heterogeneous antigen expression patterns and are known to directly suppress effector cell proliferation and persistence. T cell engineering strategies to overcome these barriers are being explored. However, strategies to simultaneously address both antigen heterogeneity and T cell longevity, while localizing anti-tumor effects at disease sites, remain limited. In this study we explore a dual antigen targeting strategy by directing independent CARs against the solid tumor targets PSCA and MUC1. To enhance functional persistence in a tumor-localized manner, we expressed the transgenic IL-7 cytokine and receptor (IL-7Rα) in respective CAR products. We now demonstrate the potency and durable antitumor effects of this binary strategy in a pancreatic tumor model.

## 1. INTRODUCTION

Immunotherapeutic intervention with chimeric antigen receptor (CAR) modified T cells has proven transformational for the treatment of hematologic malignancies such as B-cell acute lymphoblastic leukemia (B-ALL), diffuse large B-cell lymphoma (DLBCL), multiple myeloma, and non-Hodgkin’s lymphoma.^1, 2^ To date, this has led to the FDA approval of seven CAR T cell therapy products targeting antigens including CD19 and BCMA. However, efforts to extend the therapeutic benefit of CAR T cells to solid tumors have been underwhelming. Many factors contribute to this lack of efficacy including tumor heterogeneity and the lack of a ubiquitous tumor-expressed target antigen.^3, 4^ Additionally, limited T cell proliferation, persistence and functionality have been recurrent challenges impacting the anti-tumor potency of CAR engineered T cells in solid tumors.^5–7^

Several groups have applied genetic engineering strategies to address the barriers to effective CAR therapy for solid tumors. To tackle the issue of tumor immune escape due to antigen heterogeneity, combinatorial CAR targeting has been explored in preclinical models of various tumors including glioblastoma and rhabdomyosarcoma.^8, 9^ The successful clinical application of dual antigen targeting in hematologic malignancies like B-ALL (e.g., CD19 and CD22) has illustrated the feasibility of such an approach in solid tumors and is currently being investigated clinically in glioblastoma by co-targeting EGFR and IL13Rα2.^10–12^ To enhance in vivo proliferation and persistence, we and others have explored engineering approaches including the incorporation of growth-promoting cytokines such as IL-15 and conferring T cells with the ability to sequester TME-produced cytokines to drive expansion (e.g. 4/7 inverted chimeric receptor).^13–17^ We now address both barriers simultaneously by exploring the activity of a dual tumor-associated antigen (TAA) targeted approach comprised of two individual CAR T cell products targeting prostate stem cell antigen (PSCA) and Mucin1 (MUC1). To localize effector function and enhance the persistence and potency profile of the infused cells at the tumor, we further engineered CAR-PSCA T cells to produce IL-7 cytokine and the CAR-MUC1 T cells to overexpress the cognate IL-7 receptor (IL-7Rα). Here, we illustrate the benefits of this binary strategy in a pancreatic cancer model.

## 2. MATERIALS AND METHODS

### 2.1 Donor and cell lines

Peripheral blood mononuclear cells (PBMCs) were obtained from healthy donors after informed consent on protocols approved by the Baylor College of Medicine Institutional Review Board.

The cell lines CAPAN1, K562, and 293T were obtained from the American Type Culture Collection (Rockville, MD) and were grown in Dulbecco’s modified eagle medium (DMEM, GE Healthcare Life Sciences, Pittsburgh, PA) supplemented with 10% heat-inactivated fetal bovine serum (FBS) (Hyclone, Waltham, MA) and 2 mM L-GlutaMAX (Gibco BRL Life Technologies, Inc., Gaithersburg, MD). All cell lines were maintained in a humidified atmosphere containing 5% carbon dioxide (CO_2_) at 37°C.

### 2.2 Generation of retroviral constructs and retroviral supernatant

We synthesized human, codon-optimized 2^nd^ generation CAR with specificity against PSCA using the published scFv sequences,^18, 19^ which was cloned in-frame with the IgG2-CH3 domain (spacer), CD28 co-stimulation domain, and the zeta (ζ) chain of the T cell receptor (TCR) CD3 complex in an SFG retroviral backbone to make the 2nd generation CAR. We generated a 2^nd^ generation CAR with specificity against MUC1 using the published scFv sequences,^20–22^ which was cloned in-frame with the IgG2-CH3 domain (spacer), 41BB co-stimulation domain, and the ζ chain of the T cell receptor (TCR) CD3 complex in an SFG retroviral backbone. To generate the IL-7 construct, we synthesized (DNA 2.0, Menlo Park, CA) a codon-optimized sequence encoding the signal peptide and extracellular domain of human IL-7, with restriction sites *Xho*1 and *Sph1* incorporated up and downstream, respectively. To generate the IL-7Rα construct, we synthesized (DNA 2.0, Menlo Park, CA) a codon-optimized sequence encoding the signal peptide and extracellular domain of the human IL-7α, with restriction sites *Xho*1 and *Sph1* incorporated up and downstream, respectively. The IL-7 and IL-7Rα DNA inserts were incorporated into SFG retroviral vectors that contained mOrange and GFP fluorescent markers, respectively. Retroviral supernatant for these vectors was generated by transfection of 293T cells. 3×10^6^ 293T cells were plated in 10 cm tissue culture treated dishes in 10 mL Iscove’s Modified Dulbecco’s Medium (IMDM, GE Healthcare Life Sciences, Pittsburgh, PA) supplemented with 10% heat-inactivated FBS and 2 mM L-GlutaMAX. A day later, cells were transfected with PegPam, RDF, and the DNA construct loaded into GeneJuice Transfection Reagent (Millipore, Burlington, MA).^23^ Supernatant from the transfected cells were collected at 48 and 72 hrs, pooled together, and stored at-80 °C until transduction.

### 2.3 Generation of CAR T cells

To generate CAR T cells, 1×10^6^ PBMCs were plated in each well of a non-tissue culture-treated 24-well plate that had been pre-coated with OKT3 (1 mg/ml) (Ortho Biotech, Inc., Bridgewater, NJ) and CD28 (1 mg/ml) (Becton Dickinson & Co., Mountain View, CA). Cells were cultured in complete medium (RPMI-1640 containing 45% Clicks medium (Irvine Scientific, Inc., Santa Ana, CA), 10% FBS and 2 mM L-GlutaMAX), which was supplemented with recombinant human IL-2 (50 U/mL, NIH, Bethesda, VA) on day 1. On day 3 post OKT3/CD28 T blast generation, 1 mL of retroviral supernatant was added to a 24-well non-tissue culture-treated plate pre-coated with recombinant fibronectin fragment (FN CH-296; Retronectin; Takara Shuzo, Otsu, Japan) and centrifuged at 2000G for 90 minutes. OKT3/CD28 activated T cells (0.2 x 10^6^/mL) were resuspended in complete media supplemented with IL-2 (100U/mL) and then added to the wells and centrifuged at 400G for 5 minutes. To co-express IL-7 and IL-7Rα with PSCA CAR and MUC1 CAR respectively, activated T cells were transduced sequentially on days 3 and 4 – first with the CAR to generate C.P and C.M cells, and then with either IL-7 or IL-7Rα respectively to generate C.P7 and C.M7R T cells. Transduction efficiency was measured 3 days after the final transduction by flow cytometry.

### 2.4 CAPAN1 transduction

We generated CAPAN1 cell line that expressed transgenic GFP/FFLuc to facilitate bioluminescence imaging. To do this, GFP/FFLuc retroviral supernatant was plated in a non-tissue culture-treated 24-well plate (1 ml/well), which was pre-coated with a recombinant fibronectin fragment. CAPAN1 cells (0.2×10^6^/mL) were added to the plates (1 mL/well) and then transferred to a 37°C, 5% CO_2_ incubator. Transgene expression was analyzed by flow 1-week post-transduction. Cells were subsequently sorted based on GFP using a MoFlo flow cytometer (Cytomation, Fort Collins, CO).

### 2.5 Flow cytometry

The following antibodies were used in this study for T cell phenotyping: CD3-APC, CD4-Krome Orange, CD8-Pacific Blue, CD127-AF00, CD3-AF700 (Beckman Coulter Inc. Brea, CA). PSCA and MUC1 antigen expression by tumor cells was measured using anti-PSCA and anti-MUC1 (Santa Cruz Biotechnology. Inc., Dallas, TX), respectively. CAR molecules were detected using Goat anti-human F(ab’)2 antibody conjugated with AlexaFluor647 (109-606-097) (Jackson ImmunoResearch Laboratories, Inc., West Grove, PA). Cells were stained with saturating amounts of antibody (∼5uL) for 20 min at 4°C, washed (PBS, Sigma-Alrich, St. Louis, MO), and then acquired on Gallios™ Flow Cytometer (Beckman Coulter Inc., Brea, CA). Analysis was performed using Kaluza® Flow Analysis Software (Beckman Coulter Inc.).

### 2.6 ^51^Chromium-release assay

The cytotoxicity and specificity of engineered T cells was evaluated in a standard 4-6 hr ^51^Cr-release assay, using E:T ratios of 40:1, 20:1, 10:1, and 5:1. Effector T cells were co-incubated in triplicate with target cells labeled with ^51^Cr in a V-bottomed 96-well plate. At the end of the incubation period at 37°C and 5% CO_2_, supernatants were harvested, and radioactivity counted in a gamma counter. Percentage of specific lysis was calculated as follows: % specific cytotoxicity = [experimental release (cpm) – spontaneous release (cpm)]/[maximum release (cpm) – spontaneous release (cpm)] x 100.

### 2.7 T cell stimulation assays

To measure production of IL-7 by C.P7 T cells, irradiated K562 cells expressing PSCA antigen were used to stimulate C.P7 T cells at different E:T ratios. Supernatant samples were collected at different time points to measure IL-7 levels by ELISA. To evaluate growth of C.M and C.M7R T cells, 1×10^6^ irradiated CAPAN1 tumor cells were used to stimulate equal number of T cells in absence or presence of IL-7 (10ng/mL). Cells were quantified manually and using flow cytometer. Mixed T cell stimulation assay was setup by mixing 1:1 (2.5×10^5^ cells each) of C.P T cells with C.M or C.M7R T cells, and C.P7 T cells with C.M or C.M7R T cells in presence of 0.5×10^6^ irradiated CAPAN1 tumor cells. Cells were quantified periodically using flow cytometer by using CountBright™ Absolute Counting Beads (Invitrogen, Eugene, OR) and 7-AAD was added to exclude dead cells. Acquisition was halted at 2,500 beads and manufacturer’s instructions were used to calculate total cell numbers.

### 2.8 ELISA

Human Duoset ELISA kits from R&D Systems (Minneapolis, MN) were used to measure IL-7, IFN-γ, and TNF-α cytokines produced by T cells. Supernatant samples to measure IL-7 production by T cells were collected at indicated time points and stored at-20°C. IFN-γ and TNF-α were measured in supernatant obtained from T cells cultured with tumor cells overnight (∼20 hours) and stored at-20°C. All samples were thawed and processed on ice for ELISA, performed according to the manufacturer’s instructions.

### 2.9 2D co-culture experiments

For co-culture experiments, GFP/FFLuc expressing CAPAN1 tumor cells (1×10^6^ cells) were inoculated into 3D algimatrix bioscaffold (Thermo Fisher Scientific, Inc., Waltham, MA) and cultured in 6-well G-Rex devices (Wilson Wolf Manufacturing, New Brighton, MN). Three days later, 5×10^4^ C.M, 5×10^4^ C.P, C.P+C.M (2.5×10^4^ each), or C.P7+C.M7R (2.5×10^4^ each) T cells were added to tumor cells. Anti-tumor activity was monitored using the IVIS Lumina In Vivo Imaging system (Caliper Life Sciences, Hopkinton, MA) 10 minutes after adding D-luciferin (PerkinElmer, Waltham, MA) (15 mg/mL) into the culture media. Cells were then collected and recovered from the Algimatrix using Aligmatrix dissolving buffer (Thermo Fisher Scientific, Inc.). To quantify cells by flow cytometry we used CountBright™ Absolute Counting Beads (C36950; Invitrogen, Eugene, OR) and 7-AAD was added to exclude dead cells. Acquisition was halted at 2,500 beads.

### 2.10 Spheroid culture

Twelve-well tissue-culture-treated plates (Corning, NY, USA, catalog no. 353043) were coated with 200 µL of 1% (W/V) agarose (Fisher Scientific, Pittsburgh, PA, catalog no. BP160-500). 1×10^6^ Capan-1 GFP/FFLuc-labeled tumor cells were resuspended in 1 mL of T-cell medium and seeded in a final volume of 2 mL. The plates were incubated at 37°C to allow spheroid formation for 48 hours. Subsequently, the spheroids were transferred to a 6-well G-Rex (Wilson Wolf Manufacturing, New Brighton, MN, catalog no. 80240M) preconditioned with 30 mL of T-cell medium per well. Spheroid growth was monitored using IVIS Lumina In Vivo Imaging system (Caliper Life Sciences, Hopkinton, MA) 10 minutes after adding D-luciferin (PerkinElmer, Waltham, MA) (15 mg/mL) into the culture media at indicated time intervals.

### 2.11 Luciferase-based cytotoxicity assay using 3D spheroids in a G-Rex system

To test the cytotoxicity of effector cells, co-culture experiments were performed to assess the in vitro anti-tumor activity of T cells against tumor target cells. Capan-1 GFP/FFLuc-labeled tumor spheroids were generated as previously described. Two days later, they were transferred to a 6-well G-Rex in a final volume of 29 mL of T-cell medium per well. Initial bioluminescence signal produced by spheroids was measured before T cell treatment using IVIS Lumina In Vivo Imaging system 10 minutes after adding D-luciferin. GFP-or mOrange-labeled effector T cells were then added, resuspended in 1 mL of T-cell medium, resulting in a final volume of 30 mL per well. Subsequent luminescence signals were captured at the indicated intervals as described above. Cells from the wells were collected on day 14, labeled with a CD3-APC antibody (Beckman Coulter, Indianapolis, IN, catalog no. IM2467U) to distinguish GFP-or mOrange-labeled T cells, and mixed with CountBright™ Absolute Counting Beads (Invitrogen, Eugene, OR, catalog no. C36950) and 7-AAD (BD Biosciences, Franklin Lakes, NJ, catalog no. 51-68981E) to exclude dead cells. Absolute cell numbers were calculated according to CountBright™ beads manufacturer’s recommended protocol. Data analysis was performed using Kaluza® Flow Analysis Software version 2.1.

### 2.12 Statistical analysis

Data are reported as average ± standard error of mean (mean± SEM) unless stated otherwise in the figure legends. All statistical analyses were performed using GraphPad PRISM software version 10.3.1 (GraphPad Software, Inc.). Statistical differences between groups were analyzed by t-tests, one-way ANOVA, or two-way ANOVA as indicated in the figure legends. P value less than 0.05 was regarded as statistically significant.

## 3. RESULTS

### 3.1 Limited T cell persistence restricts durable tumor control in a dual target setting

To target PSCA and MUC1, both TAAs known to be frequently expressed in pancreatic tumors,^24, 25^ we generated two human codon-optimized second generation CARs with specificity for PSCA (C.P) and MUC1 (C.M), containing CD28 and 41BB co-stimulatory endodomains, respectively, which were fused to the CD3ζ activation motif (**Fig. 1A**).^26, 27^ Retroviral transduction led to efficient expression of both CARs in healthy donor-derived primary human T cells, as illustrated in **Fig. 1B**. Next, using a standard 5-hr ^51^Cr-release assay, we evaluated the cytolytic activity of these CAR-modified cells against CAPAN1 pancreatic cancer cells, which express both target TAAs, albeit at different levels (PSCA – 95.1% and MUC1 – 68.5%; **Fig. 1C**). CAPAN1 cells were killed by both the C.P (**Fig. 1D**) and C.M (**Fig. 1E**) while demonstrating minimal reactivity against antigen negative 293T cells. As anticipated, superior killing was associated with higher target antigen expression levels. To next assess the long-term anti-tumor activity of these T cells, either alone or in combination, we utilized a previously established extended in vitro co-culture system.^27^ GFP/FFLuc-modified CAPAN1 tumor cells (CAPAN1.GFP/FFLuc) embedded in Algimatrix were cultured with C.P, C.M, or both (C.P:C.M; 1:1) in 6-well G-Rex devices, and monitored by bioluminescence imaging. As shown in **Fig. 1F**, untreated tumor cells grew unimpeded, while treatment with individual CAR products each demonstrated some level of tumor control. However, co-administration of both C.P and C.M T cells at 1:1 ratio with total (C. P+C.M) T cell numbers equal to single-treated conditions, demonstrated superior anti-tumor effects. Nevertheless, tumor regrowth was eventually detected in all conditions. We confirmed that this was not due to antigen loss by evaluating PSCA and MUC1 surface levels on outgrowing tumor cells (on day 20), which demonstrated expression of both antigens comparable to the untreated condition (**Fig. 1G**). Interestingly, a second administration of T cells (on day 27) resulted in a decline in tumor bioluminescence (**Fig. 1H**). This outcome illustrates the targetability of the residual tumor cells and implicates limited T cell persistence as the likely cause of tumor regrowth. Taken together, these results demonstrate that targeting more than one TAA in heterogeneous tumors improves overall tumor killing. However, suboptimal T cell persistence limits this benefit, resulting in a lack of durable anti-tumor responses.

**Figure 1:**
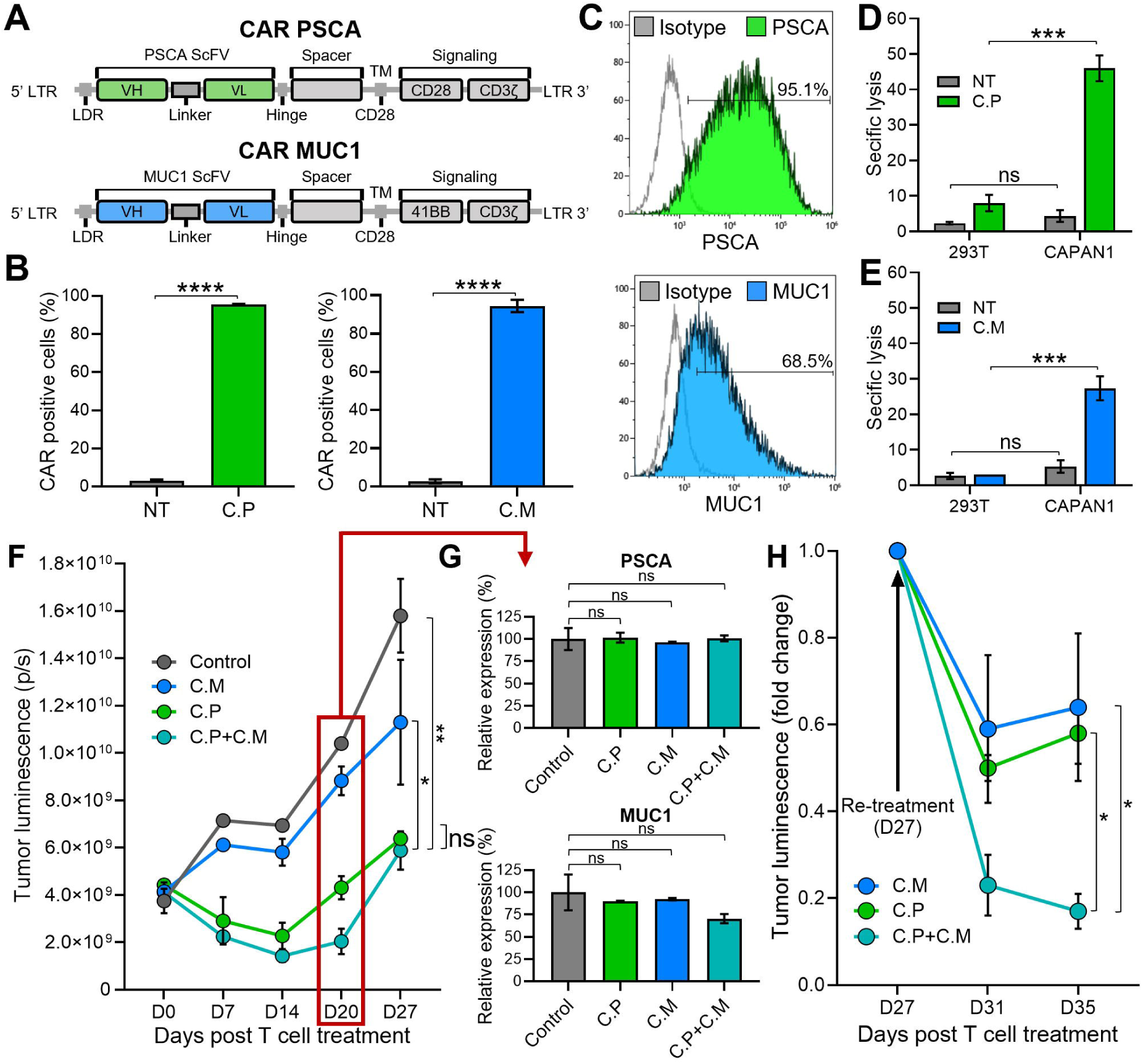
Limited T cell persistence restricts durable tumor control in a dual target setting **(1A)** Illustrations of the CARs targeting PSCA (top) and MUC1 (bottom) used to generate CAR T cells. **(1B)** PSCA (left) and MUC1 (right) CAR expression in C.P and C.M T cells measured by flow cytometry 3 days post-transduction. Non-transduced (NT) T cells used as negative controls (t-tests, n=3, ns = no significant difference, *p<.05, **p<.01, ***p<.001, ****p<.0001). **(1C)** Detection of PSCA (top) and MUC1 (bottom) expression by CAPAN1 tumor cells measured using flow cytometry. **(1D)** Cytolytic activity of C.P T cells against 293T (PSCA negative control) and CAPAN1 tumor cells measured using Cr-release assay at the effector to target ratio (E:T) of 20:1. NT cells used as controls (t-tests, n=3, ns = no significant difference, *p<.05, **p<.01, ***p<.001, ****p<.0001). **(1E)** Anti-tumor activity of C.M T cells against 293T and CAPAN1 cells measured using Cr-release assay at the effector to target ratio (E:T) of 20:1 (t-tests, n=3, ns = no significant difference, *p<.05, **p<.01, ***p<.001, ****p<.0001). **(1F)** CAPAN1 tumor luminescence measured by IVIS imaging in a long-term co-culture assay with CAR T cells in G-Rex 6-well plates. Untreated (tumor-only) condition used as control (one-way Anova on day 27, n=3, ns = no significant difference, *p<.05, **p<.01, ***p<.001, ****p<.0001). **(1G)** PSCA (top) and MUC1 (bottom) expression by CAPAN1 cells from the co-culture experiment in (1F), measured on day 20 by flow cytometry. All T cell treated conditions normalized to the control (t-tests, n=3, ns = no significant difference, *p<.05, **p<.01, ***p<.001, ****p<.0001). **(1H)** CAPAN1 luminescence in C.M, C.P, and C.P+C.M treated conditions in (1F) after re-treatment with respective CAR T cells on day 27 (t-tests on day 35, n=3, ns = no significant difference, *p<.05, **p<.01, ***p<.001, ****p<.0001).

### 3.2 Engineering CAR T cells to secrete and efficiently utilize IL-7 cytokine

Potent and sustained T cell activation, persistence and anti-tumor activity requires three stimulatory signals (signal 1 – antigen recognition, signal 2 - co-stimulation, and signal 3 - cytokine).^28^ While our C.P and C.M cells receive signals 1 (CD3ζ) and 2 (CD28 or 41BB co-stimulatory domains, respectively) through the CAR, they lack signal 3. Thus, we explored whether the incorporation of transgenic cytokine support would promote superior anti-tumor effects. We evaluated the potential of various cytokines including IL-2 and IL-15, both of which have been shown to effectively promote T cell expansion.^29, 30^ However, IL-2 and IL-15 can be utilized by inhibitory cells including tumor-infiltrating regulatory T cells, which are known to suppress the anti-tumor function of infused CAR T cells.^31–34^ Furthermore, systemic administration of these cytokines can cause abnormalities such as capillary leak syndrome, hypotension, fever, and vomiting due to systemic toxicity affecting multiple organ systems.^35, 36^ To balance the need to support T cell expansion without promoting toxic side effects, we therefore designed a binary T cell strategy in which C.P cells were modified to secrete IL-7 (C.P7) (a homeostatic cytokine that augments the proliferation and survival of T cells, particularly naïve and memory subsets), and C.M cells to overexpress the IL-7Rα (C.M7R).

To test the feasibility of this binary system, we first transduced C.P cells with a retroviral vector encoding the human IL-7 gene linked to mOrange fluorescent protein by an IRES sequence (**Fig. 2A**, top). T cells were efficiently transduced to express both CAR and IL-7 transgenes (**Fig. 2A**, bottom, representative donor), with an average of >85% double-transduced cells (**Fig. 2B**). As measured by ELISA, antigen (K562 cells engineered to express PSCA) stimulated C.P7 T cells secreted IL-7 (**Fig. 2C**), which progressively increased with increasing antigen exposure (**Fig. S1A**) and duration of culture (**Fig. S1B**). We confirmed that C.P7 T cells retained their target antigen specificity and ability to produce effector cytokines. In a standard chromium release assay, C.P7 T cells were able to selectively kill PSCA-expressing targets (**Fig. S1C**) and exhibited a cytokine profile that was comparable to C.P cells (**Fig. S1D)**. Taken together, our results demonstrate that C.P cells can be engineered to efficiently secrete IL-7, which is produced in an antigen stimulation-dependent manner without compromising the antigen specificity or anti-tumor function of the CAR.

**Figure 2:**
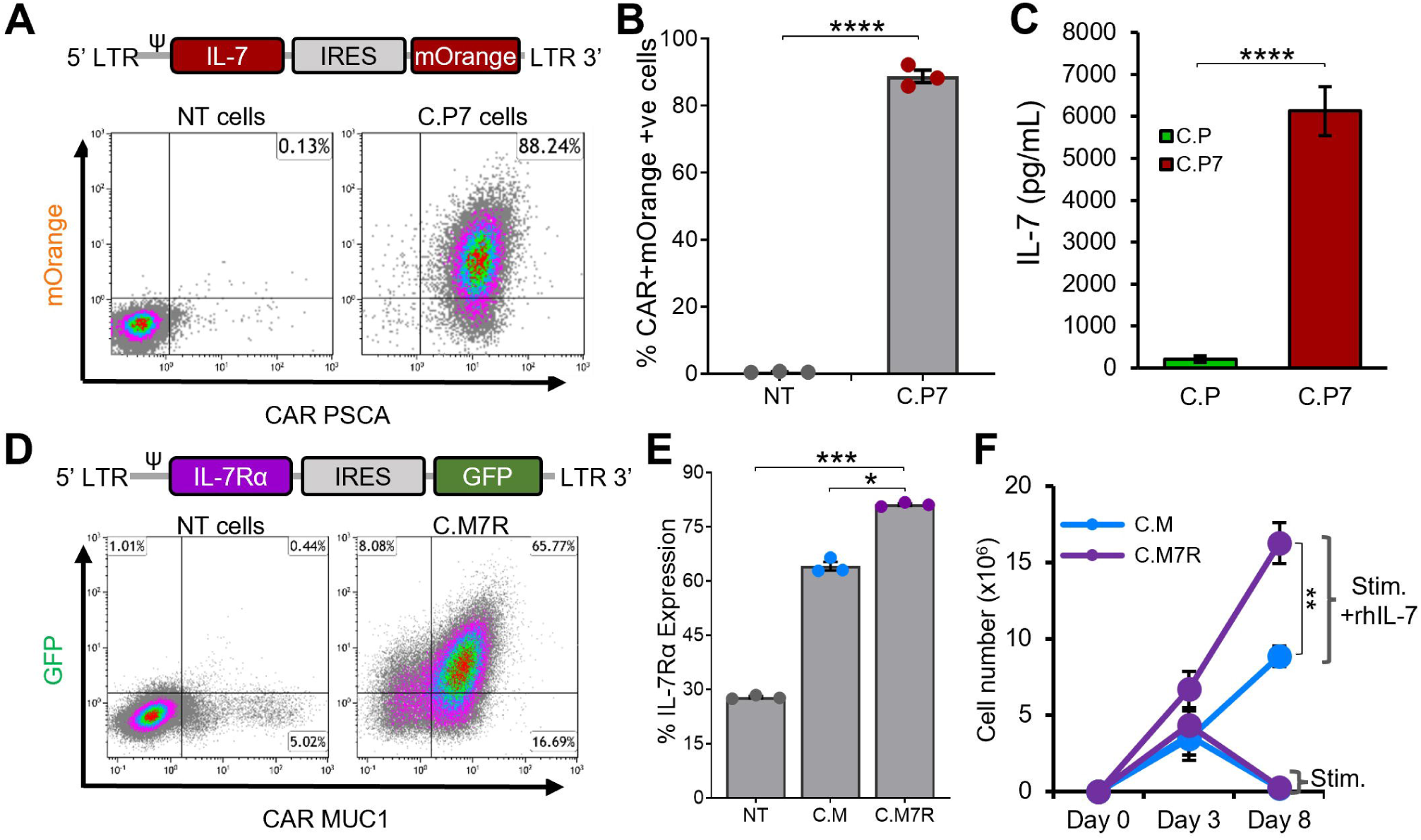
Engineering CAR T cells to secrete and efficiently utilize IL-7 cytokine **(2A)** Illustrations of the IL-7 cytokine construct containing mOrange fluorescent protein for transgene detection by flow cytometry (top) and detection of both the PSCA CAR and IL-7 transgene expression in C.P7 T cells from a representative donor after serial transduction (bottom). **(2B)** Summary data indicating PSCA CAR and IL-7 double-transduced T cells compared to NT cells (t-tests, n=3, ns = no significant difference, *p<.05, **p<.01, ***p<.001, ****p<.0001). **(2C)** Production of IL-7 by C.P7 cells after stimulation with irradiated K562 cells engineered to express PSCA, measuring using ELISA (t-tests, n=3, ns = no significant difference, *p<.05, **p<.01, ***p<.001, ****p<.0001). **(2D)** Diagram illustrating the IL-7Rα construct containing GFP for transgene detection (top) and flow cytometry data for a representative donor demonstrating the expression of both the MUC1 CAR and the IL-7Rα transgenes in C.M7R T cells. **(2E)** Summary data comparing IL-7Rα detection by flow cytometry in NT, C.M, and C.M7R cells (one-way ANOVA, n=3, ns = no significant difference, *p<.05, **p<.01, ***p<.001, ****p<.0001). **(2F)** Quantification of C.M and C.M7R T cells using trypan blue exclusion during culture with irradiated CAPAN1 tumor cells in presence or absence of recombinant IL-7 cytokine (t-tests on day 8, n=3, ns = no significant difference, *p<.05, **p<.01, ***p<.001, ****p<.0001).

While both chains of the IL-7R complex – IL-7Rα (CD127) and common γ chain (CD132) are endogenously expressed by lymphocytes, including T cells, native IL-7Rα expression is known to be downregulated upon exposure to IL-7.^37, 38^ Additionally, antigenic stimulation has also been reported to reduce IL-7Rα expression by T cells in a non-IL-7 dependent manner.^39^ We explored the regulation of endogenous IL-7Rα in our C.M T cells and, consistent with previous findings, confirmed that upon IL-7 cytokine or antigen stimulation, the receptor was downregulated (**Fig. S2A)**. To enhance the capacity of C.M cells to utilize C.P7-produced IL-7 (**Fig. 2A-C** and **S1**), we transduced C.M cells with a retroviral vector encoding human IL-7Rα linked to GFP by an IRES sequence (**Fig. 2D**, top). This resulted in the production of dual-transgenic T cells (**Fig. 2D**, bottom) with superior IL-7Rα expression compared to C.M cells (**Fig. 2E**). Notably, this receptor overexpression resulted in greater expansion of C.M7R T cells compared to C.M cells when stimulated with irradiated CAPAN1 tumor cells in the presence of recombinant IL-7 (**Fig. 2F**), consistent with enhanced IL-7 utilization.

Finally, to ensure that overexpression of IL-7Rα did not negatively impact the functional capacity of our CAR T cells, we assessed antigen-specificity, effector cytokine production, and responsiveness to other common γ chain family cytokines. As illustrated in **Fig. S2B** and **S2C**, the antigen specificity and cytolytic function of C.M7R T cells was similar to that of C.M T cells, as was the secretion of effector cytokines IFN-γ and TNF-α upon exposure to the tumor cells. Furthermore, C.M7R cells retained their ability to respond to other common γ chain family cytokines such as IL-2 and IL-15, illustrated by the similar growth pattern of C.M and C.M7R T cells (**Fig. S2D-E**). In summary, these results demonstrate that engineering C.M cells to overexpress IL-7Rα improves their ability to utilize IL-7 without adversely impacting their antigen specificity, effector cytokine production capacity, and ability to utilize other common γ chain cytokines such as IL-2 and IL-15.

### 3.3 IL-7 engineered C.P7 T cells support the expansion of C.M7R overexpressing IL-7Rα

We next evaluated whether the combination of C.P7 and C.M7R cells supported superior T cell expansion. First, we explored whether the quantity of IL-7 produced by C.P7 cells following antigen exposure was sufficient to drive T cell expansion. To assess this, we cultured C.M7R cells in conditioned medium harvested 48 hours after stimulating C.P7 cells or C.P (control lacking IL-7) with irradiated CAPAN1 tumor cells at E:T of 1:1. As shown in **Fig. 3A**, expansion of C.M7R cells in C.P7 conditioned medium was significantly greater than that seen in cytokine-free medium or C.P conditioned medium. Next, to assess if antigen-activated C.P7 cells were able to support C.M7R cell expansion, we co-cultured both T cell populations with irradiated CAPAN1 tumor cells (E:E:T ratio of 1:1:2) and evaluated cell outgrowth by flow cytometry. As illustrated in **Fig. 3B**, co-stimulating C.M or C.M7R cells with non-cytokine-producing C.P T cells (green lines) resulted in only short-term expansion followed by rapid T cell contraction during the second week of culture. In contrast, co-stimulation of C.M or C.M7R cells with C.P7 T cells produced a marked increase in total T cell numbers (red lines), with superior output seen in the binary condition (solid red line). Further in-depth analysis of the binary condition and separation of C.P7 from C.M7R cells based on GFP (surrogate for transgenic IL-7Rα) expression revealed the presence of both cell types, though the C.M7R population, consistent with their ability to utilize C.P7 produced cytokine (**Fig. 3C**).

**Figure 3:**
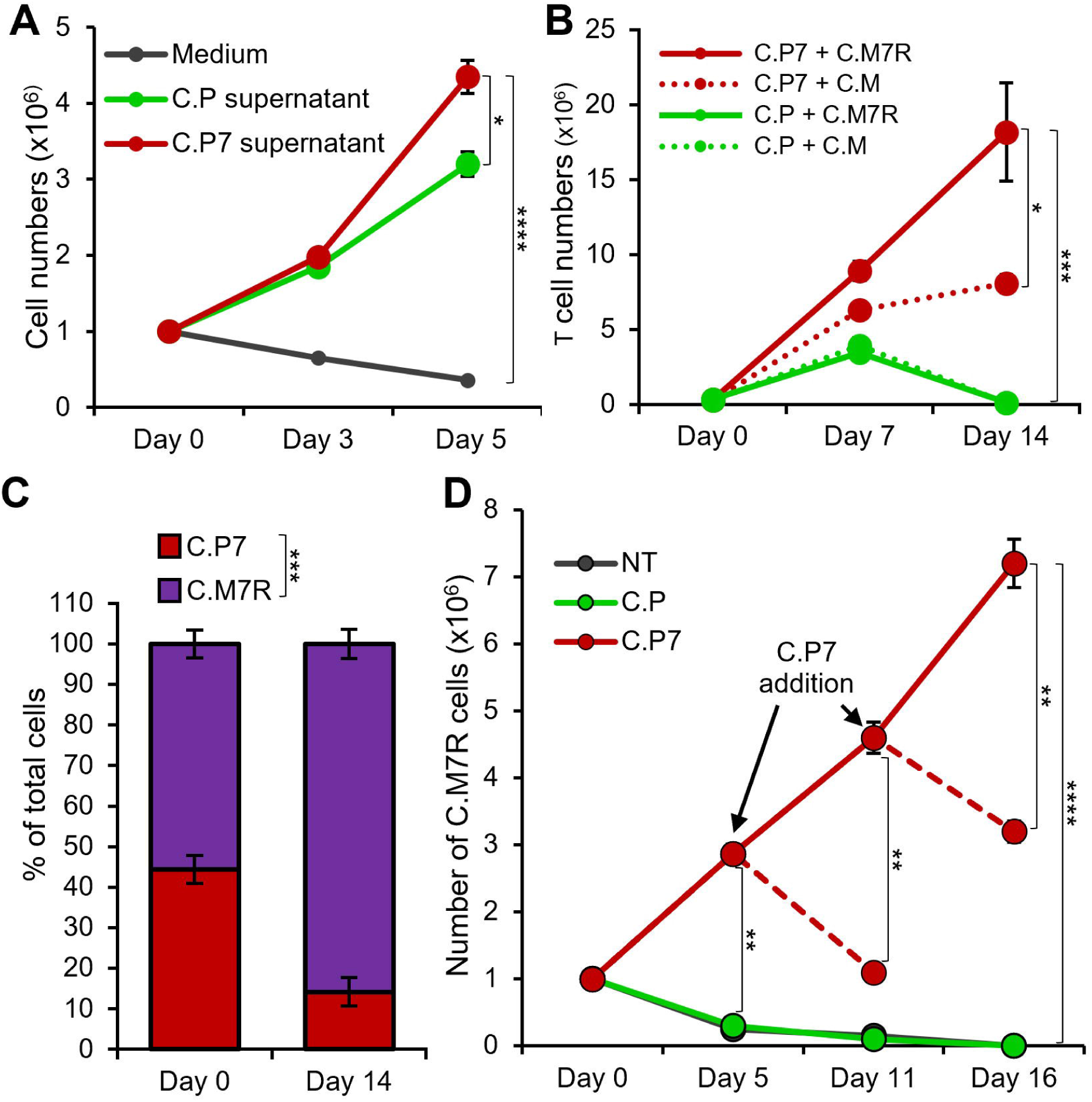
IL-7 engineered C.P7 T cells support the expansion of C.M7R overexpressing IL-7R**α (3A)** Growth of C.M7R T cells in cell culture medium, and conditioned medium collected from C.P or C.P7 T cells stimulated with irradiated CAPAN1 tumor cells. Cells were quantified by manual counting (t-tests on day 5, n=3, ns = no significant difference, *p<.05, **p<.01, ***p<.001, ****p<.0001). **(3B)** Total cell numbers in irradiated CAPAN1-stimulation co-cultures of the indicated T cell types (t-tests on day 14, n=3, ns = no significant difference, *p<.05, **p<.01, ***p<.001, ****p<.0001). **(3C)** Differentiation of C.P7 and C.M7R T cells based on GFP (surrogate for transgenic IL-7Rα) using flow cytometry to quantify the proportion of these cells in the C.P7+C.M7R condition at the start and the end of the stimulation experiment in (3B) (t-tests on day 14, n=3, ns = no significant difference, *p<.05, **p<.01, ***p<.001, ****p<.0001). **(3D)** Flow cytometric quantification of C.M7R T cells cultured with NT (control), C.P, or C.P7 T cells pre-stimulated with irradiated CAPAN1. Arrows indicate time points at which additional pre-stimulated C.P7 T cells were added. Dashed lines indicate conditions which were monitored for an additional time point without the additional dose of pre-stimulated C.P7 T cells (t-tests at indicated time-points, n=3, ns = no significant difference, *p<.05, **p<.01, ***p<.001, ****p<.0001).

One potential advantage of this binary T cell approach is the ability to control and support proliferation and persistence of the receptor-expressing population, with the strategic delivery of the cytokine-producing cell component. To test whether repeated administration of C.P7 would indeed support this outcome, we cultured C.M7R cells with either non-transduced (NT) control T cells, C.P, or C.P7 T cells stimulated with irradiated CAPAN1 tumor cells for 48 hours at the E:T of 1:1 and monitored the expansion profile of C.M7R cells. As shown in **Fig. 3D**, we observed robust C.M7R T cell expansion on day 5 post co-culture with activated C.P7 cells, while limited or no expansion was detected when co-cultured with NT or C.P cells. Furthermore, periodic addition of activated C.P7 T cells (every 5-6 days) resulted in a boosting effect, producing an increase in C.M7R cell numbers, whereas a single administration of stimulated C.P7 cells resulted in an eventual decrease in the number of C.M7R T cells. In summary, these results demonstrate that transgenic IL-7 produced by C.P7 cells supports T cell expansion, particularly of cells engineered to overexpress IL-7Rα. Furthermore, strategic administration of C.P7 cells can be used to prolong the expansion and persistence of C.M7R cells.

### 3.4 Dual-targeted binary T cells exhibit enhanced anti-tumor activity and persistence

To evaluate if the binary strategy results in enhanced and durable anti-tumor effects, we compared the tumor killing ability of single (C.P or C.M, individually), dual (C.P+C.M), and the binary T cells (C.P7+C.M7R) in the long-term co-culture assay with CAPAN1 tumor cells described in **Fig. 1**. An initial decrease in tumor luminescence was observed in all treatment conditions, indicating tumor-killing by CAR T cells (**Fig. 4A**, representative tumor wells and **4B**, summarized results). However, the tumors eventually outgrew by day 20 (**Fig. 4B**) in the C.P, C.M, and C.P+C.M treatment conditions as seen previously (**Fig. 1F**). In contrast, the C.P7+C.M7R T cell combination produced superior anti-tumor effects with no detectable tumor outgrowth (**Fig. 4A, B**). This enhanced anti-tumor activity, as anticipated, coincided with increased total T cell expansion/persistence in the binary treatment group (**Fig. 4C**).

**Figure 4:**
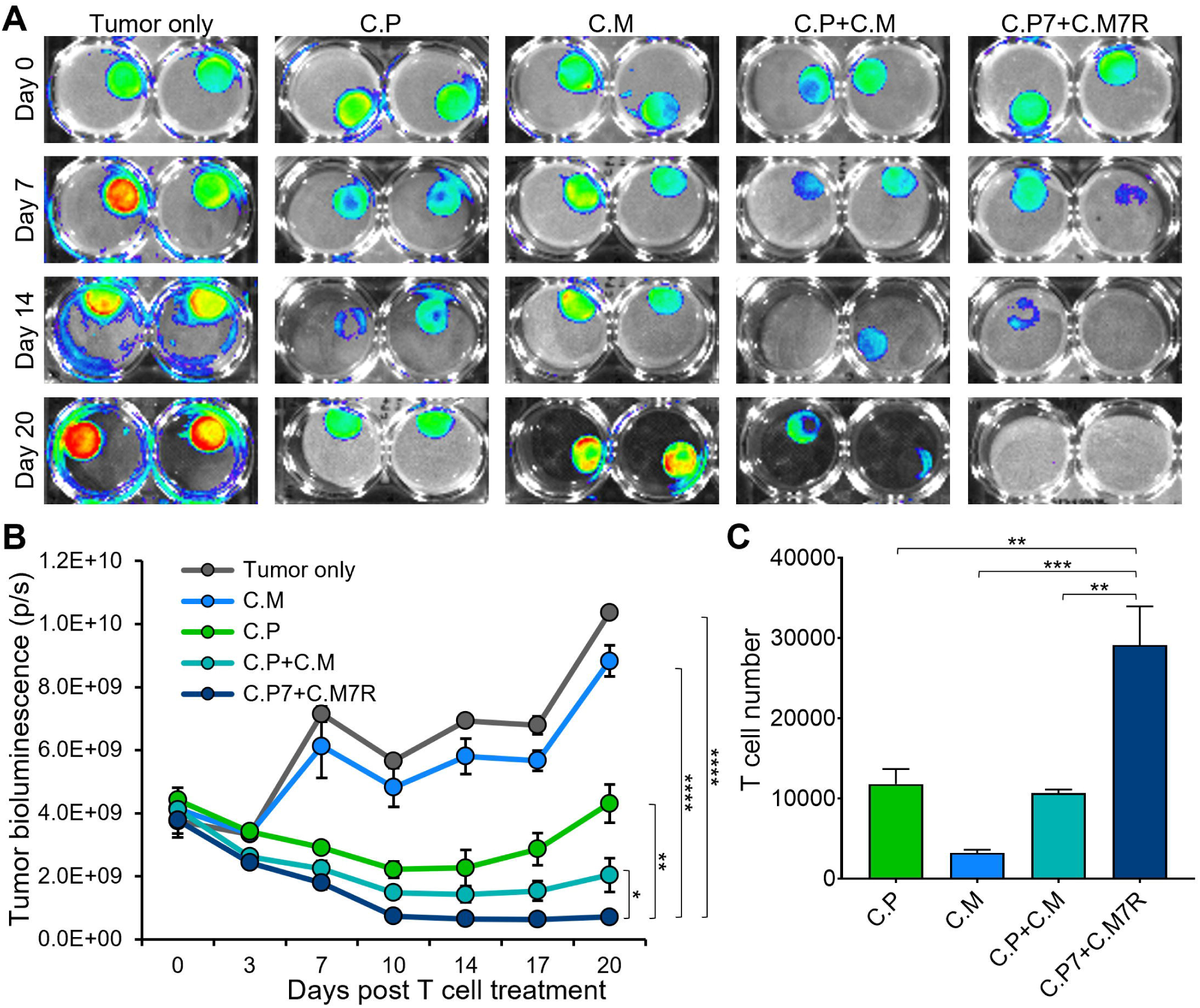
Dual-targeted binary T cells exhibit enhanced anti-tumor activity and persistence **(4A)** Representative bioluminescent images of GFP/FFLuc expressing CAPAN1-seeded Algimatrix cultures in the long-term co-culture assay using 6-well G-Rex devices. Two replicates shown for each condition **(4B)** Quantification of bioluminescence signal from the CAPAN-1 cells [illustrated in (4A)] during the co-culture experiment under indicated treatment conditions (t-tests on day 20, n=3, ns = no significant difference, *p<.05, **p<.01, ***p<.001, ****p<.0001). **(4C)** Live T cell counts using flow cytometry on day 20 for the different treatment conditions in (4B) (one-way ANOVA, n=3, ns = no significant difference, *p<.05, **p<.01, ***p<.001, ****p<.0001).

### 3.5 Binary T cells demonstrate superior anti-tumor activity against pancreatic spheroids

3D culture systems more closely replicate *in vivo* characteristics with tumor spheroids providing a model that better mimics the solid tumor architecture and physiology.^40^ Hence, we next explored the benefits of our binary strategy with pancreatic cancer cell-derived 3D tumor spheroids. We first established spheroids from GFP/FFLuc-labeled CAPAN1 tumor cells and optimized the growth conditions in 6-well G-Rex devices to ensure spheroid viability for an extended period of time (**Fig. S3A-B)**. To confirm that CAPAN1 TAA expression was maintained in 3D cultures, we isolated tumor cells at two different time points (day 4 and 25) after spheroid formation and measured the expression of PSCA and MUC1, both of which were maintained throughout the culture period of 25 days (**Fig. S3C**). To assess anti-tumor activity we next co-cultured tumor spheroids alone (Tumor only), and with C.P+C.M, or C.P7+C.M7R T cells at the calculated E:T (based on initial number of tumor cells used to generate spheroids) of 1:40. Bioluminescence imaging of the spheroids demonstrated a decline in tumor burden in both T cell treated conditions (**Fig. 5A**, representative wells and **Fig 5B**, summarized results). However, the anti-tumor response in the C.P+C.M treated conditions, as observed in the previous co-cultures (**Fig. 4**), was less robust compared to the C.P7+C.M7R treatment conditions (**Fig. 5A** and **5B**). To confirm if this enhanced tumor control in the binary condition was due to improved T cell expansion/persistence, we isolated and quantified T cells from the co-cultures on day 14. Similar to our observations in the Algimatrix co-culture system (**Fig. 4C**), higher numbers of total T cells were quantified in the C.P7+C.M7R combination compared to C.P+C.M treatment (**Fig. 5C**). In summary, these data demonstrate the binary system we developed to augment CAR T cell activity against heterogeneous tumors is effective against the physiologically more relevant 3D spheroids and confirm the results of our previous co-culture assessments.

**Figure 5:**
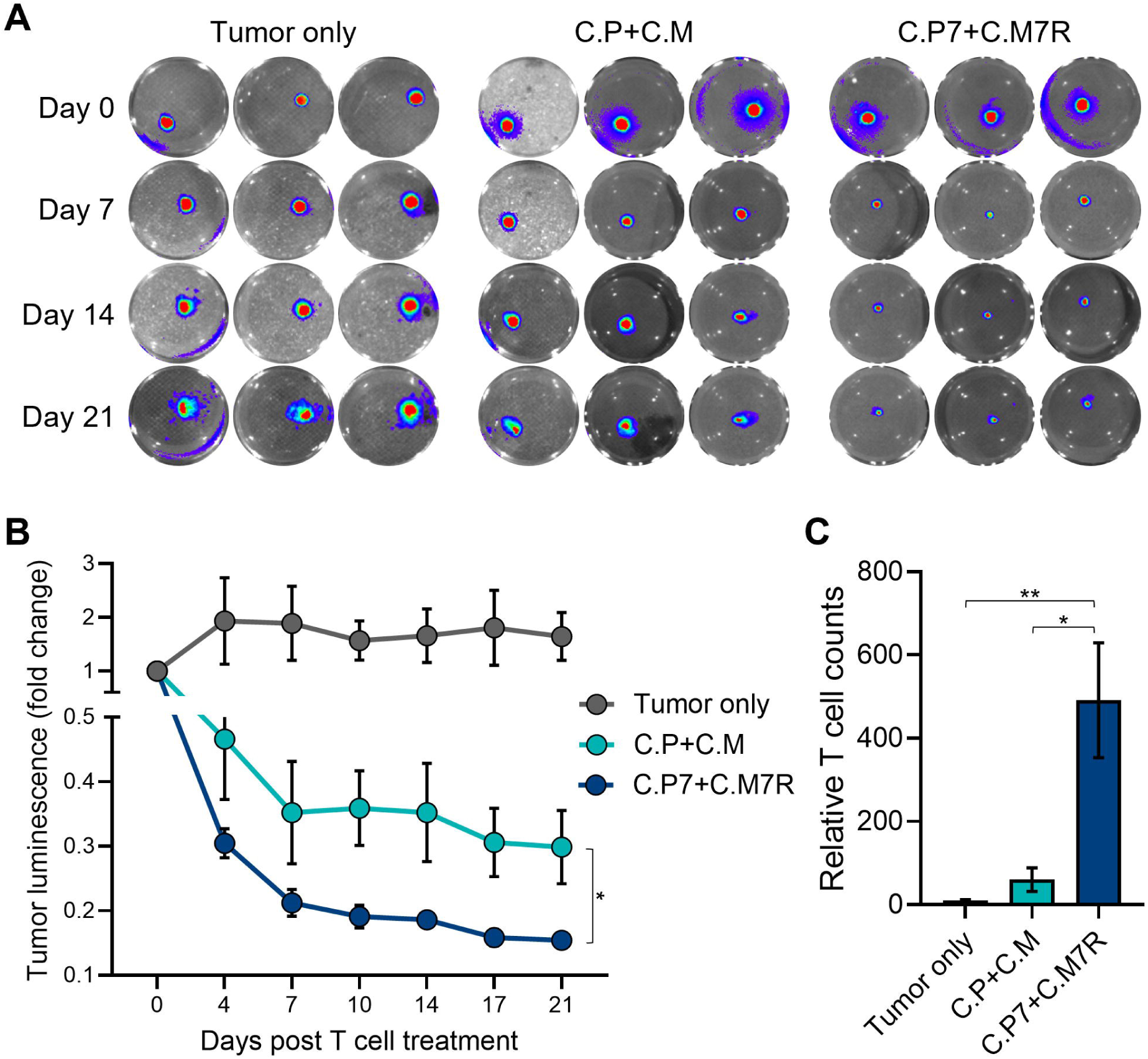
Binary T cells demonstrate superior anti-tumor activity against pancreatic spheroids **(5A)** Representative bioluminescent images of GFP/FFLuc expressing CAPAN1 tumor cells generated spheroids during long-term co-culture. Three replicates illustrated for each condition. **(5B)** Quantitate bioluminescence data for the CAPAN1 spheroids in absence of treatment or during co-culture with indicated T cell types (t-tests on day 21, n=9, ns = no significant difference, *p<.05, **p<.01, ***p<.001, ****p<.0001). **(5C)** Live T cell counts for the different treatment conditions in (5B) obtained using flow cytometry on day 14 (t-tests, n=4, ns = no significant difference, *p<.05, **p<.01, ***p<.001, ****p<.0001).

## 4. DISCUSSION

In this study, we explored the feasibility of implementing a binary T cell strategy to simultaneously address two barriers to successful immunotherapy for solid tumors – namely suboptimal anti-tumor activity due to antigen heterogeneity and limited expansion and persistence of tumor-challenged CAR T cells. To address the issue of heterogeneity, we generated two CAR T cell products with specificities for the TAAs PSCA and MUC1, and to enhance T cell proliferation and persistence, we further engineered our CAR-PSCA T cells to produce IL-7 cytokine and CAR-MUC1 T cells to overexpress the IL-7Rα. We reasoned that the provision of this tumor localized cytokine support would promote the expansion and persistence of the transferred cells in a safe and regulated manner, given the co-dependence of the two cellular products. Indeed, using 2D and 3D pancreatic cancer models, we demonstrated the superiority of our binary approach over single-pronged strategies, with enhanced T cell expansion and persistence, resulting in more pronounced and sustained anti-tumor responses. Although the current study focused on pancreatic cancer models, the binary platform can be adapted and applied to enhance the efficacy of a range of tumor-targeted products such as TCR-modified T cells and CAR-NK cells.

Solid tumors typically exhibit heterogeneous antigen expression patterns, which creates the risk of escape from mono-targeted cell therapies. One strategy to prevent this outcome is by targeting two or more tumor antigens simultaneously, either using T cells engineered to co-express multiple CARs or by co-administering two or more mono-targeted T cell products.^41, 42^ This approach has been explored clinically to address antigen-negative B-ALL relapses, which occur in 10-20% of CD19 CAR T cell recipients, by simultaneously targeting a second antigen like CD20 or CD22.^11, 43^ For example, CD19/22 dual-targeted CAR T cells demonstrated superior complete remission rates (98%, n=50/51, NCT03614858) compared to CD19-only targeted therapy (83%, n=122/147, NCT03919240).^44, 45^ This strategy has been explored in the preclinical models of solid tumors like glioblastoma (HER2 and IL13Rα2 CARs) and rhabdomyosarcoma (FGFR4 and B7-H3 CARs) and shown to result in superior durability of anti-tumor effects.^8, 9^ Additionally, an ongoing phase I study investigating bivalent CAR (targeting EGFR and IL13Rα2) engineered T cells recently reported transient reductions of recurrent glioblastoma, which has median overall survival of less than a year, in all treated patients (n=6).^12^ These results demonstrate the potential of dual-TAA targeting in aggressive tumors with poor prognosis; however, the poor proliferation and persistence of the infused CAR T cells within solid tumors remains a hurdle.

In solid tumors, inadequate in vivo T cell expansion and persistence, which ultimately diminishes anti-tumor effects, have remained a persistent challenge.^3–5, 42^ This limited T cell proliferation and durability of the infused T cells can be due to the hostile tumor microenvironment of solid tumors as well as T cell intrinsic factors such as suboptimal CAR design and T cell exhaustion.^3, 46, 47^ To augment T cell proliferation and durability of effect, a variety of strategies have been explored including engineering T cells to produce proliferation/persistence promoting cytokines like IL-2, IL-15, and IL-12.^48, 49^ For instance, Hombach and colleagues demonstrated that co-administration of mesenchymal stem cells (MSCs) genetically engineered to secrete IL-7 and IL-12 significantly improved the anti-tumor activity of CEA-targeted CAR T cells in a preclinical model of colon cancer.^50^ Shum et al applied a different strategy to enhance longevity but limiting the effects to specific cells. They engineered GD2 CAR T cells to express a synthetic IL-7R (C7R), which is constitutively active due to cysteine/proline insertions in the transmembrane region that activate STAT5 resulting in IL-7 stimulation in absence of the ligand. Subsequently, in a neuroblastoma model these investigators demonstrated enhanced tumor-localized proliferation and anti-tumor activity of GD2-CAR-C7R compared with CAR alone.^51^ However, it should be noted that this feature of constitutive signaling/activation is also seen in some pre-T cell acute leukemias, and hence incorporation of the C7R molecule in conjunction with a safety switch is advisable.^52^ In contrast, the approach we explored has the benefit of being “self-regulating” while efficiently delivering cytokine at the tumor site as – (i) IL-7 production is dependent on antigenic stimulation of C.P7 T cells, and this (ii) allows tumor localized C.M7R cells to benefit. Furthermore, the segregated expression of cytokine and receptor in different cell populations minimizes the risk of uncontrolled proliferation due to autonomous cytokine signaling.

While conventional preclinical paradigms typically rely on in vivo animal studies in immune-compromised mice engrafted with tumor xenografts, we chose to implement a novel 3D tumor spheroid model system, which our group developed to enable the long-term assessment of effector proliferation, persistence and CAR potency. We and others have found the expression of TAAs, including PSCA and MUC1, to be similar between 3D models and tumor xenografts, and tumor spheroids present a bulk mass that recapitulates the solid tumor architecture and tumor-produced extracellular matrix, which is well suited for initial preclinical screening of complex therapeutics, as assessed in the current study.^40, 53, 54^ Of note, our implementation of such a platform aligns well with recent FDA guidance advocating for the replacement of animal testing with effective, human-relevant methods that can serve to expedite the preclinical evaluation of novel therapeutics.^55^

In conclusion, our study illustrates that providing tumor-localized IL-7 cytokine support via T cells engineered to secrete as well as utilize the cytokine in a dual-antigen targeting binary system enhances the expansion and anti-tumor function of CAR T cells. Importantly, cytokine production and T cell expansion was detected only in the presence of tumor cells expressing the targeted antigens. These results illustrate the potential of our binary strategy to improve the efficacy of CAR T cells in solid tumors by overcoming antigen heterogeneity and limited T cell expansion/persistence.

## 6. AUTHOR CONTRIBUTIONS

AGTC: Investigation, Methodology, Formal analysis, Writing – review & editing; MKM: Methodology, Writing – review & editing, AG: Investigation, Writing – review & editing; ND: Investigation, Writing – review & editing; JV: Conceptualization, Methodology, Funding acquisition, Writing – review & editing; AML: Conceptualization, Supervision, Resources, Funding acquisition, Writing – review & editing; PB: Conceptualization, Investigation, Formal analysis, Methodology, Supervision, Writing – original draft.

## 8. ACKNOWLEDGEMENTS

We are thankful to the Texas Children’s Hospital Small Animal Imaging Facility and the shared resources support from the Dan L. Duncan Comprehensive Cancer Center. We also thank: Dr.

Malcolm Brenner and Dr. Norihiro Watanabe for their advice on the study, and Walter Mejia for the illustrations and artwork used in the manuscript.

## 9. DATA AVAILABILITY STATEMENT

The data generated in this study are included in the manuscript figures and the Supplementary Material. Additional/raw data are available from the corresponding author upon reasonable request.

