## Supplementary data for "IL-7 armed binary CAR T cell strategy to augment potency against solid tumors"

### Supplementary Figure 1

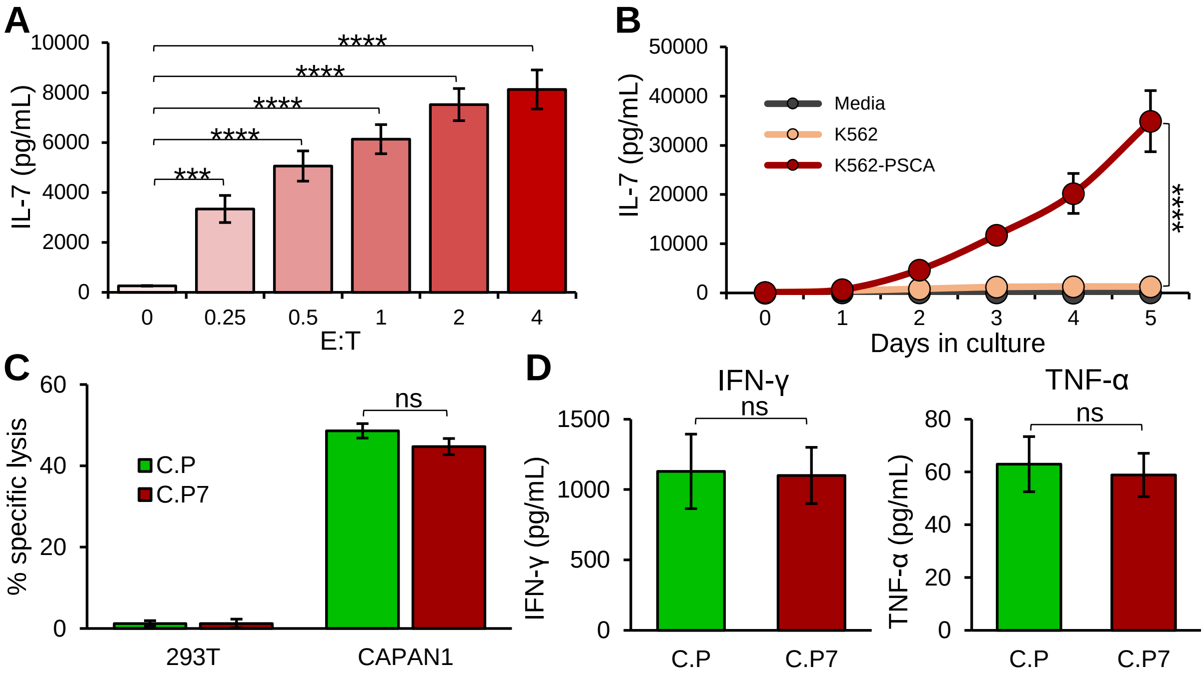


**Supplementary Figure 1: (S1A)** IL-7 production by C.P7 T cells measuring using ELISA when stimulated with increasing number of irradiated K562 cells engineered to express PSCA (one-way ANOVA, n=3, *p<.05, **p<.01, ***p<.001, ****p<.0001). **(S1B)** IL-7 detection in supernatants obtained at indicated time points from C.P7 T cells cultured in medium alone, with irradiated K562 cells (PSCA negative), or with irradiated K562 cells expressing PSCA (t-tests on day 5, n=3, *p<.05, **p<.01, ***p<.001, ****p<.0001). **(S1C)** Anti-tumor activity of C.P and C.P7 T cells against 293T and CAPAN1 cells measured using $^{51}$Cr-release assay at the effector to target ratio (E:T) of 20:1 (t-tests, n=3, ns = no significant difference, *p<.05, **p<.01, ***p<.001, ****p<.0001). **(S1D)** Detection of IFN-γ and TNF-α cytokines by ELISA in the supernatants obtained from C.P and C.P7 T cells after overnight culture with CAPAN1 tumor cells (t-tests, n=3, ns = no significant difference, *p<.05, **p<.01, ***p<.001, ****p<.0001).

### Supplementary Figure 2

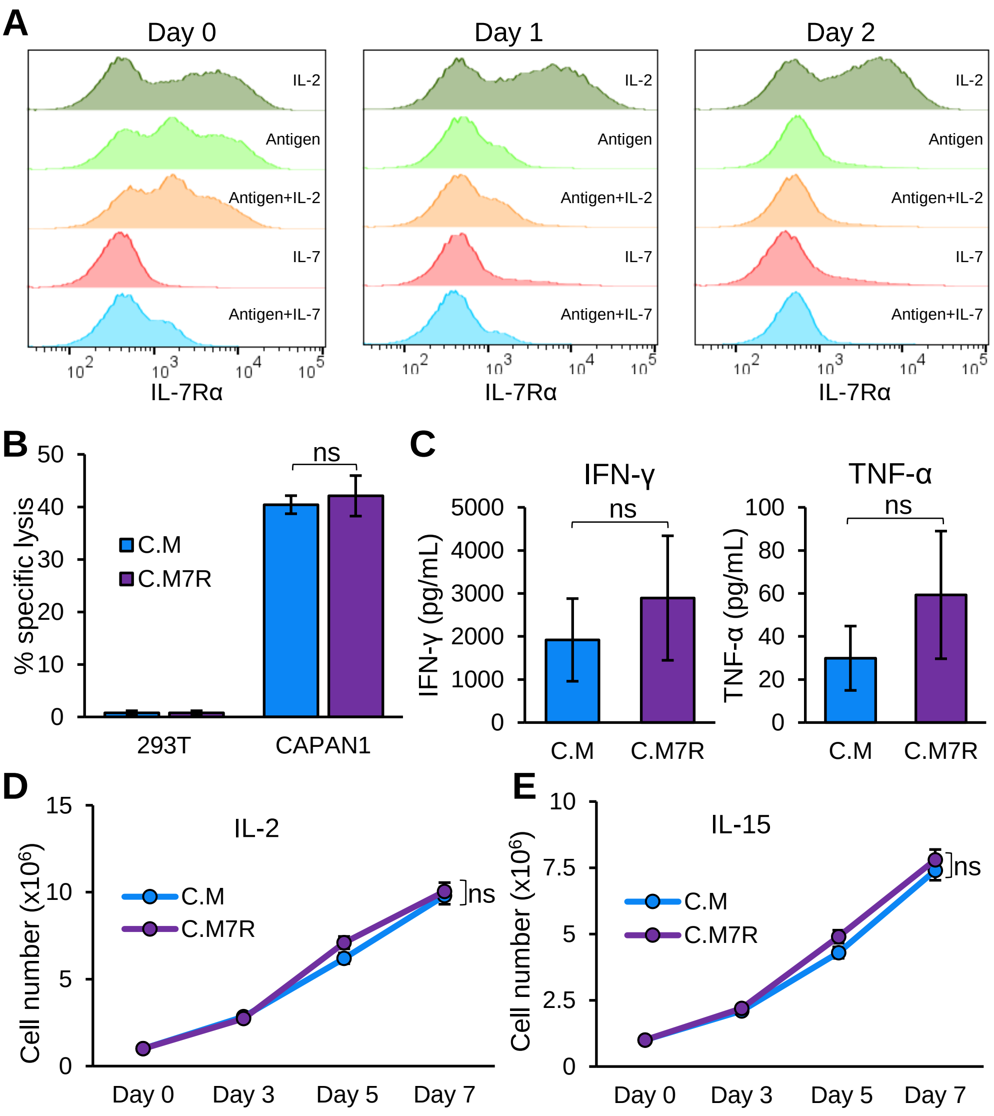


**Supplementary Figure 2: (S2A)** Flow cytometric detection of IL-7Rα on the surface of C.M T cells under the indicated culture conditions **(S2B)** Anti-tumor activity of C.M and C.M7R cells against 293T and CAPAN1 cells measured using $^{51}$Cr-release assay at the E:T of 20:1 (t-tests, n=3, ns = no significant difference, *p<.05, **p<.01, ***p<.001, ****p<.0001). **(S2C)** IFN-γ and TNF-α production by C.M and C.M7R T cells after overnight culture with CAPAN1 tumor cells, measured by ELISA (t-tests, n=3, ns = no significant difference, *p<.05, **p<.01, ***p<.001, ****p<.0001). **(S2D)** Expansion of C.M and C.M7R T cells in culture medium supplemented with IL-2 cytokine, cell counts obtained by manual (trypan blue exclusion) counting (t-tests on day 7, n=3, ns = no significant difference, *p<.05, **p<.01, ***p<.001, ****p<.0001). **(S2E)** Expansion of C.M and C.M7R cells in IL-15 cytokine supplemented culture medium, cell counts obtained by manual counting (t-tests on day 7, n=3, ns = no significant difference, *p<.05, **p<.01, ***p<.001, ****p<.0001).

### Supplementary Figure 3

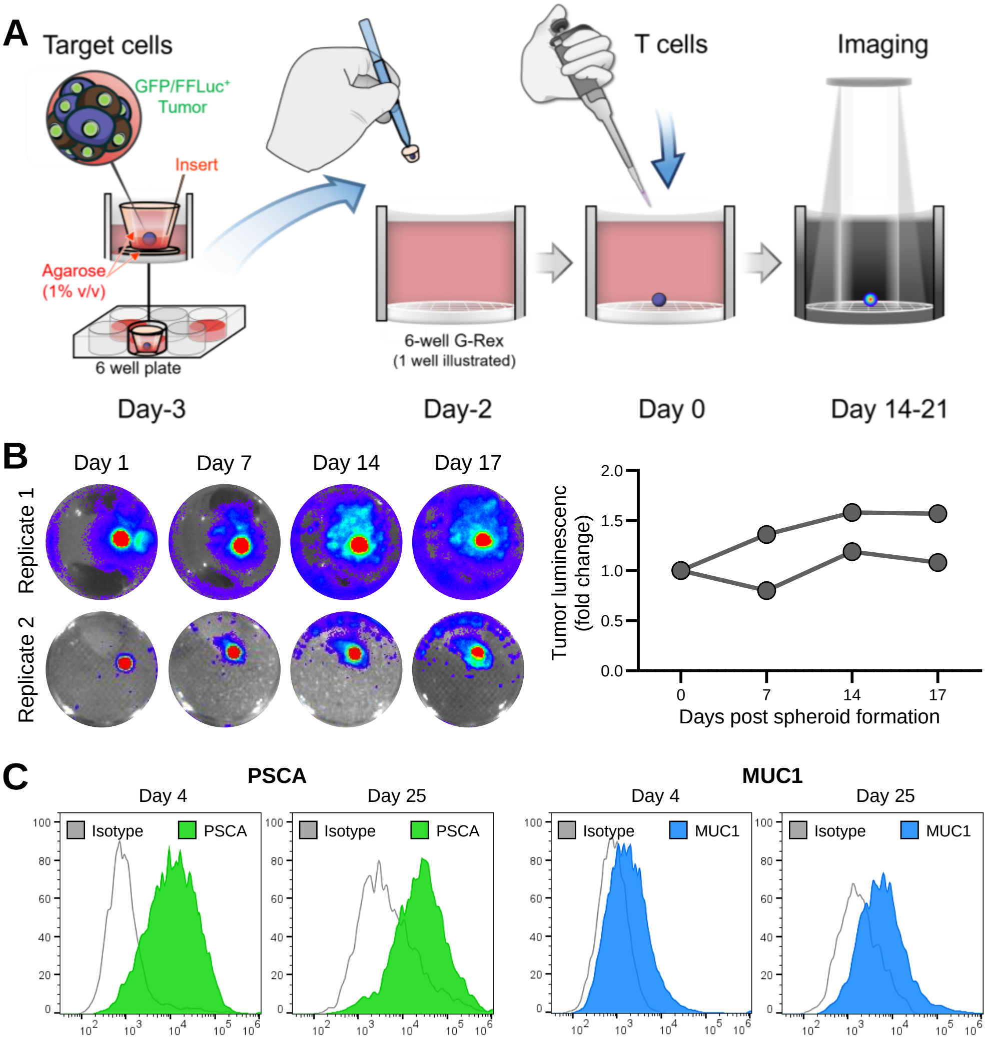


**Supplementary Figure 3: (S3A)** Illustration of pancreatic spheroids generation process in G-Rex. (**S3B**) Representative bioluminescent images (left) and time-course fold change in luminescence (right) of GFP/FFLuc expressing CAPAN1 tumor cells generated spheroids cultured in the 6-well G-Rex device. Three replicates shown. **(S3C)** Flow cytometric detection of PSCA (left) and MUC1 (right) antigens by tumor cells in the spheroids measured at an early (day 4) and late (day 25) time point after spheroid formation.
